# Context Shapes Alpha-Band Electrophysiological Signatures under Psilocybin

**DOI:** 10.64898/2026.10.06.757155

**Authors:** Vidushani Dhanawansa, Devon Stoliker, Leonardo Novelli, Mana Biabani, Adeel Razi, Levin Kuhlmann

## Abstract

**Background:** Psilocybin is a serotonergic psychedelic that provides a pharmacological tool for examining neural activity associated with altered states of consciousness. Existing studies report reduced power (4-30 Hz) in posterior regions and altered functional connectivity (FC) at rest, but have typically involved small samples and coarse anatomical parcellations. Additionally, perceptual engagement has been shown to attenuate psilocybin-related whole-brain FC changes. However, how naturalistic context modifies psilocybin’s frequency-specific electrophysiological effects remains incompletely characterised.

**Methods:** We analysed the largest psilocybin electroencephalography (EEG) dataset to date (n=56), with matched baseline recordings acquired before psilocybin administration. EEG was recorded during movie viewing, eyes-closed rest, guided meditation, and music listening. Cortical source activity was estimated using minimum-norm estimation and analysed across six canonical frequency bands (1-80 Hz). Spectral power, alpha peak frequency (APF), and magnitude-squared coherence within and between seven functional networks were quantified. Psilocybin effects were assessed using permutation testing with FDR correction, and linear mixed-effects models separated the contributions of drug, context, and their interaction to alpha-band connectivity.

**Results:** At rest, psilocybin decreased alpha (8-13 Hz) power, increased gamma-range power, and elevated APF. Alpha-band FC decreased within and between higher-order networks implicated in top-down control. Relative to eyes-closed rest, psilocybin-context interactions showed that structured sensory environments attenuated or reversed psilocybin-related alpha-band connectivity changes.

**Conclusions:** These findings identify alpha-band activity as a key electrophysiological correlate of psilocybin-related changes in cortical communication. Structured sensory input reshapes psilocybin-related cortical dynamics such as alpha-band connectivity across large-scale cortical networks, with the strength and direction of this modulation differing by context.

## 1 Introduction

Agonism of the 5-HT_2*A*_ receptor by serotonergic psychedelics [44] can induce altered states of consciousness, positioning these compounds as valuable tools for investigating the neural mechanisms of perception, cognition and conscious experience. The emerging therapeutic potential of psychedelics such as psilocybin (the principal psychoactive alkaloid in Psilocybe mushrooms) further motivates investigation of their neural mechanisms [10, 25, 31, 32]. However, psychedelic effects are shaped by both internal state and external context, raising the question of whether context also modifies their neural expression. Heightened environmental sensitivity is a central characteristic of psychedelic experiences [16], and many clinical trials reporting promising outcomes for psychological distress, mood disorders, and addiction have placed particular emphasis on ‘setting’, including manipulation of the therapeutic environment [15].

fMRI studies have shown that psilocybin alters large-scale functional connectivity in a network-, metric-, time-, and context-dependent manner[27, 52, 57]. Under perceptual tasks (simple auditory-visual matching task), psilocybin-related whole-brain FC changes are attenuated[57]. In addition, psilocybin reduces the difference in global functional connectivity (GFC) between eyes-open and eyes-closed states (including resting-state, music listening and meditation)[63]. However, electroencephalography (EEG) and magnetoencephalography (MEG) offer a crucial complement to the strengths of fMRI by resolving frequency-specific neural activity at millisecond timescales. This capability enables frequency-resolved analysis of neural dynamics, including alpha- and gamma-band activity, providing insight into distinct aspects of cortical organisation[33]. Furthermore, these modalities provide a more direct measure of neural activity that are not dependent on the neurovascular coupling underlying fMRI blood oxygen level–dependent (BOLD) signals. This may be particularly advantageous in psychedelic research, given that serotonergic modulation can alter cerebral vascular tone and cerebral blood flow, potentially complicating the interpretation of haemodynamic measures [20, 43].

Prior M/EEG studies have investigated psychedelic-induced changes in power spectral density (PSD), signal complexity and FC under the resting-state[10]. Reductions in lower frequency oscillatory power under psychedelics including N,N-Dimethyltryptamine(DMT)[67, 68] and psilocybin[38, 39, 43], have been consistently reported, particularly within the parietal and occipital regions[4, 49]. However, with the exception of Kometer et al. [38], existing studies administering a single full psychedelic dose have relied on cohorts of fewer than 20 participants, limiting statistical power and generalisability [29].

Barnett et. al.[4] investigated the frequency specific directed and undirected FC in cortical and sub-cortical regions using the same cohort as [43], reporting reduced connectivity within coarse anatomical groupings, including frontal, parietal, and sensorimotor regions, in the alpha and beta bands. Similarly, Kometer et. al. quantified the FC between the Brodmann areas which correlated with aspects of the psychedelic experience[38]. However, substantial scope remains to characterise psilocybin-related FC within and between functionally defined cortical networks, such as the default mode, which allows a more direct systems-level framework for interpreting large-scale brain organisation than coarse anatomical parcellations. To date, psilocybin-related FC modulation of functionally relevant networks across multiple frequency bands and naturalistic contexts has not been comprehensively characterised.

Increasing efforts have been made to quantify frequency-specific neural effects of psychedelics under different naturalistic conditions[41, 55, 56]. Complementary MEG-LSD studies show that greater external stimulation was associated with attenuated LSD-induced changes in whole-brain signal complexity and alpha-band power, alongside context-dependent restructuring of canonical networks in the alpha and beta bands[41, 56]. These findings suggest a competitive interaction between psychedelic effects and sensory context. However, these findings derive from a single MEG study of LSD with relatively small cohorts (n=14–20), leaving it unclear whether the same competitive interaction between psilocybin and sensory context is evident in EEG-derived spectral power, and functional connectivity within and between key canonical networks, and whether it generalises across a broader set of naturalistic conditions in a sufficiently powered sample.

To address these gaps in understanding, we analysed EEG recordings from 56 healthy participants with no prior psychedelic experience across resting state, mindfulness meditation, music listening, and passive movie viewing conditions, each paired with matched no-psilocybin baselines. We estimated cortical source activity and quantified spectral power and functional connectivity within canonical brain networks across a broad frequency range. Following initial analyses identifying prominent and context-dependent alpha-band connectivity changes, linear mixed-effects models were used to separate the contributions of psilocybin, context, and their interaction within this band.

## 2 Methods

### 2.1 Participants and Experimental Procedures

The dataset was acquired at Monash Biomedical Imaging (MBI) and is part of a more comprehensive psychedelic imaging dataset, comprising fMRI and EEG data from sixty-five psychedelic-naive healthy participants. Participants aged 18–55 (37.3 *±* 10.7; 30 female, 35 male). The present study focuses exclusively on an expanded analysis of the EEG data; see Stoliker et al. for the corresponding imaging findings[63]. The study protocol was approved by the Monash University Human Research Ethics Committee. The trial was registered with the Australian New Zealand Clinical Trials Registry under the registration number ACTRN12621001375842. The research was performed in accordance with all relevant guidelines and regulations, and written informed consent was obtained from all participants.

Baseline (no-psilocybin) scanning took place approximately one week prior to psilocybin administration. Recordings included four conditions: movie of clouds moving (no-sound, 5 min), resting state (5 min), guided meditation (5 min), and music listening (7 min). All participants then received a fixed single oral dose of 19mg of psilocybin, with EEG recording beginning approximately 200 minutes post-administration: participants were taken for an MRI session (not reported here) approximately 80 minutes post-dosing, 75 minutes of MRI acquisition, and an approximately 20-40 minutes of EEG setup time. Psilocybin’s half-life is roughly 300-400 minutes, so most participants were still experiencing significant subject effects of the drug. Analyses relied on within-subject contrasts; therefore, only participants with complete data in both sessions were included (movie: n=53, other: n=56), with exclusions due to technical failure or incomplete recording.. Prior to discharge, participants completed a range of self-report measures (Figure 1). See the data descriptor for further details [47].

**Fig. 1:**
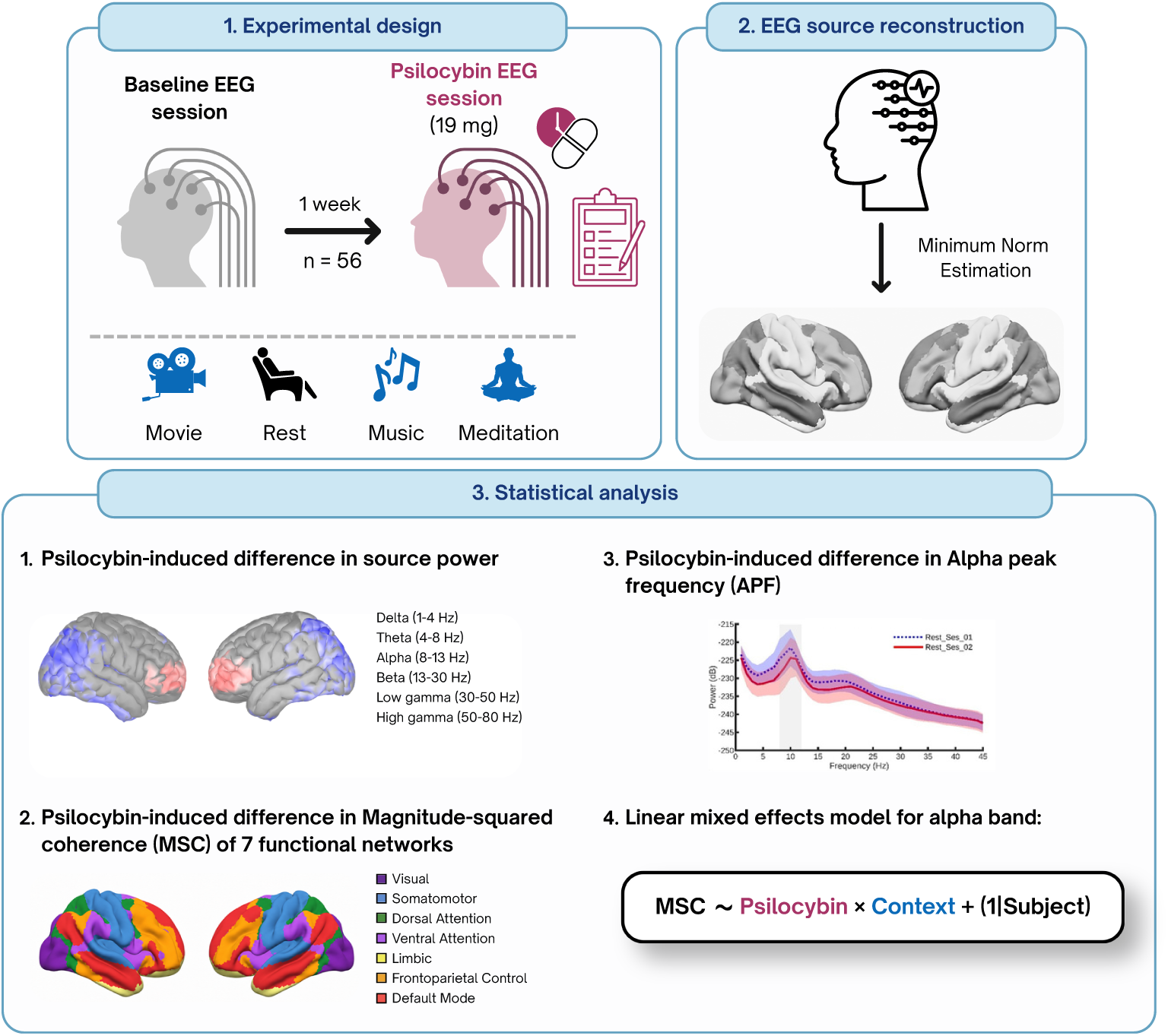
Overview of the experimental design and EEG analysis workflow. Participants underwent a baseline EEG session followed by a second EEG session approximately one week later after administration of 19 mg psilocybin. Each session included recordings across movie-watching, resting state, music listening, and meditation. EEG data were source reconstructed using minimum norm estimation, and psilocybin-related differences were examined through source-level spectral power, alpha peak frequency, and magnitude-squared coherence across seven functional networks. Linear mixed-effects models were used to assess drug, context, and drug-by-context effects.

### 2.2 EEG Acquisition and Preprocessing

The EEG was acquired from 64 electrodes using an actiCAP slim cap, positioned according to the standardized 10-10 system[19]. Signals were recorded using a BrainAmp MR Plus amplifier with BrainVision Recorder software (version 1.22.0001), all from Brain Products, GmbH, Germany. The FCz electrode served as the reference and the FPz electrode served as the ground for online recording, with electrode impedances maintained below 10 kΩ. EEG was sampled at 500 Hz with a 0.01–1000 Hz bandpass filter. The raw data were acquired as a single continuous recording over all conditions and then split into four separate files using the event markers for the start and end of each condition.

EEG preprocessing followed the RELAX pipeline (v1.1.3)[2] implemented in MATLAB (R2024b) using functions from EEGLAB (v2021.0)[22] and FieldTrip[48] toolboxes. The data was band-pass filtered (0.25-80 Hz) using a fourth-order zero-phase Butterworth filter, with a notch filter (47-53 Hz) to filter out line noise. Following the RELAX pipeline, bad electrodes were initially rejected using the PREP pipeline[11], and a multi-channel weiner filter (MWF)[59] was applied to reduce muscle, blink and horizontal eye movement artefacts. Data was re-referenced to the robust average reference. Remaining artifactual components were identified and rejected using ICLabel and wavelet enhancing ICA[51]. Electrodes rejected during the cleaning process were then restored through spherical interpolation. Finally, all preprocessed data were visually inspected for data quality.

### 2.3 EEG source reconstruction

Cortical source reconstruction was performed using minimum-norm estimation (MNE). In this framework, the source current distribution is estimated under a cortical source constraint and a Gaussian prior on source amplitudes with a specified source covariance matrix[3]. Electrode positions from the acticAP template were co-registered to individual anatomy using the nasion and left and right pre-auricular fiducials. As AFz and Iz channels were not represented in the electrode-position template, these channels were excluded from source-level analyses. The individual head models were generated using a Boundary Element Model (BEM) generated with 1922 vertices per layer for the scalp, outer skull, and inner skull layers. Default relative conductivities for the scalp, skull and brain layers were set to 1, 0.0125 and 1, respectively[65].

For the MNE based source reconstruction, the noise covariance matrix was computed from 0-20s segment of rest baseline data and only the sensor variance estimates (diagonals of covariance matrix) were retained. The source estimation resulted in 15002 dipoles constrained to the cortical surface. The source orientation remained constrained normal to the cortical surface and depth weighting was applied to reduce the superficial-source bias of MNE. Source reconstruction was performed in MATLAB (R2024b) using Brainstorm (v 3.250811)[65] and OpenMEEG (v 2.4.1) [30].

### 2.4 Source-level spectral power analysis

The Welch periodogram was computed with 2sec Hann-windowed epochs and 50% overlap, generated from source-level data segments from 20s onwards for the baseline and psilocybin recordings of all participants. The power within the following frequency bands were extracted: Delta (1-4 Hz), Theta (4-8 Hz), Alpha (8-13 Hz), Beta (13-30 Hz), Low gamma (30-50 Hz), High gamma(50-80 Hz). To facilitate group analysis, the source power maps were first projected to the ICBM152 template cortical surface in Brainstorm and then smoothed by applying Gaussian kernel of 3-mm full width at half maximum (FWHM). Statistical group analysis was performed on each frequency band where a paired t-test between the power maps of the baseline and psilocybin session was computed. 5000 permutations were computed for each test. t-images were thresholded at p*<*0.05 and Benjamini-Hochberg correction for multiple comparisons across across all bands and contexts was applied at a False Discovery Rate (FDR) of 0.05[9]. Furthermore, given previous evidence that alpha peak frequency (APF) is modulated under LSD[14], we computed APF for five regional clusters (frontal, occipital, parietal, temporal, and cingulate) defined using the Desikan–Killiany atlas [23].

### 2.5 Functional Connectivity

The functional connectivity (FC), was inferred through the magnitude-squared coherence (MSC) between the time-series of the source reconstructed regions[26]. MSC was selected to quantify frequency-specific functional coupling across cortical networks and has been applied in EEG studies of depression, providing clinical precedent[40].

The MSC between time series *x*(*t*) and *y*(*t*) is defined as

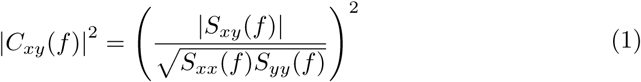

where *S_xy_*(*f*) is the cross-spectral estimate and *S_xx_*(*f*) and *S_yy_*(*f*) are the auto-spectra of signals *x*(*t*) and *y*(*t*), respectively.

Functional connectivity was quantified within and between seven canonical networks defined by the Yeo7 atlas [70], using the 100-region Schaefer cortical parcellation. The principal components of signals in each parcel was extracted, band-pass filtered into the predefined frequency bands and transformed using the Hilbert transform. The cross-spectrum and corresponding auto-spectra were averaged across all time samples within each condition before normalisation to obtain the MSC. Mean MSC within and between network pairs was compared between baseline and psilocybin sessions using permutation-based Wilcoxon signed-rank tests (2-sided at p*<*0.05, 5000 permutations and FDR correction at 0.05).

To further explore the influence of psilocybin, context, and their interaction on network connectivity, exploratory linear mixed-effects (LME) analyses were conducted within the alpha band, selected based on the preceding non-parametric analysis showing significant between-session differences and differential psilocybin effects across contextual conditions (see Supplementary Material for details):

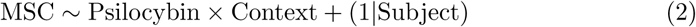

Two sets of models were constructed with different connectivity outcomes: (1) MSC of each network pair and (2) mean MSC within each network (e.g. vis–vis) and between each network and all other networks (e.g. vis–other). FDR correction was applied across all tested model parameters at *q <* 0.05.

We then computed the spearman correlations between the psychometric scales recorded during the psilocybin session and psilocybin-related: (1) within-network FC changes and (2) mean FC changes between each network and the rest of the networks, both within the resting-state. The chosen scales were the 11-Dimensional Altered States of Consciousness (11D-ASC), Mystical Experience Questionnaire (MEQ) and Ego-Dissolution Inventory (EDI) [6, 24, 46, 64] (see Supplementary material for details).

## 3 Results

This study characterised source-level EEG changes induced by psilocybin across rest and naturalistic contexts. Results are presented for: spectral power and alpha peak frequency, functional connectivity within and between canonical cortical networks, and linear mixed-effects modelling of context-dependent alpha-band connectivity.

### 3.1 Spectral analysis of source reconstructed EEG

The statistically significant differences of spectral power of the psilocybin vs baseline session are visualized in the source maps in Figure 2 and Table S1. Under the resting state, in comparison to the baseline state, psilocybin was linked to (at p*<*0.05, FDR corrected):

- An increase in the frontal delta power and decrease in the occipital delta power.
- A widespread strong decrease of theta band power.
- A decrease of alpha power in key regions of the DMN and visual networks, including the precuneus, posterior cingulate, paracentral lobule, cuneus, insula, fusiform gyrus, inferior parietal lobule and parahippocampal gyrus.
- A widespread increase in power in the gamma bands.

**Fig. 2:**
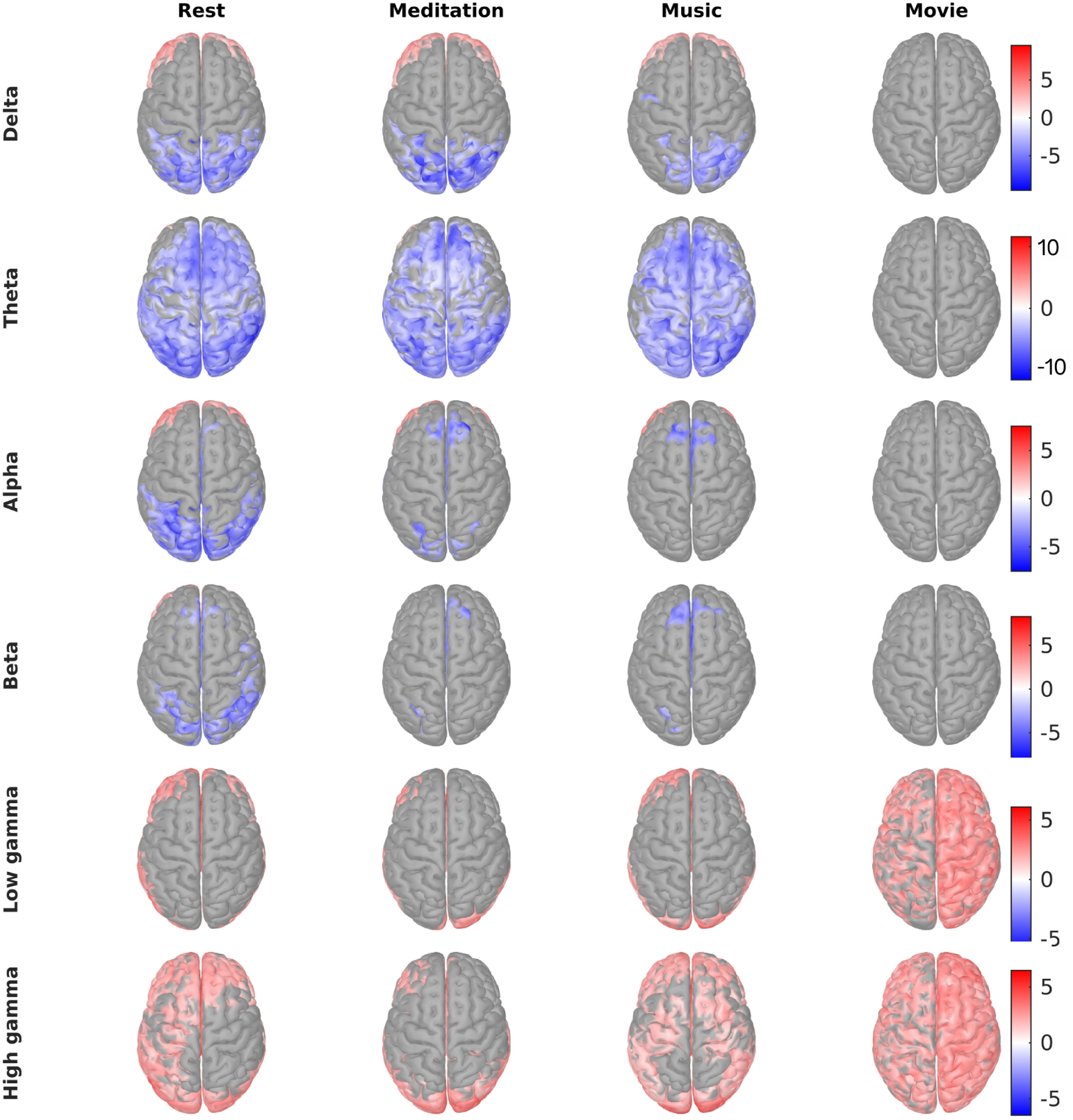
Statistical parametric maps of significant (p*<*0.05, corrected) psilocybin-related changes in source spectral power relative to the baseline condition under resting state, meditation, music and movie-watching. Positive t-values indicate a psilocybin-related increase in power.

Under the eyes-closed meditation and music conditions, psilocybin was associated with similarly pronounced reductions in delta and theta power across broadly overlapping cortical regions as observed during resting state. In contrast, in the alpha band, significant parietal power reductions observed during resting state, overlapping with default mode and visual network regions, were not observed in the other conditions (Figure S1). In the movie condition, only the psilocybin-related increase in gamma power survived FDR correction. A paired t-test revealed that psilocybin resulted in a significant (p*<*0.05) increase in alpha peak frequency (APF) across most region clusters under each context (Table 1).

**Table 1:** t-values of paired t-test testing an increase in alpha peak frequency during the psilocybin session relative to baseline across cortical region clusters (**p *<* 0.005; *p *<* 0.05).

| Region | Rest | Meditation | Music | Movie |
| --- | --- | --- | --- | --- |
| Frontal | 4.056** | 2.649* | 2.966** | 3.463** |
| Occipital | 1.588 | 2.925** | 1.784* | 2.383* |
| Parietal | 3.300** | 1.272 | 2.669** | 2.556* |
| Temporal | 1.992* | 2.169* | 3.778** | 2.906** |
| Cingulate | 3.582** | 2.855** | 3.207** | 1.098 |

### 3.2 Functional Connectivity

#### 3.2.1 Resting state functional connectivity (RSFC)

In the alpha band, psilocybin was linked to decreased RSFC (p*<*0.05, FDR corrected) in all network pairs except within the visual and limbic network (Figure 3 and Figure S2). RSFC increased in the gamma bands with significant increases occurring across most network pairs in the high gamma band. psilocybin-related RSFC changes in low- to mid-frequency bands (1-30 Hz) within and between key networks implicated in top-down control positively correlated with ratings of audio-visual synesthesia (AVS)(r*>*0.2, p*<*0.05, not FDR corrected). This included the control (or frontoparietal), default mode, salience/ventral attention (S/VAN), dorsal attention (DAN) and somatomotor (SMN) networks. Ratings of changed meaning of percepts and impaired control and cognition correlated with within- and between-network RSFC of the S/VAN, DAN and SMN, across different frequencies (Figure 4).

**Fig. 3:**
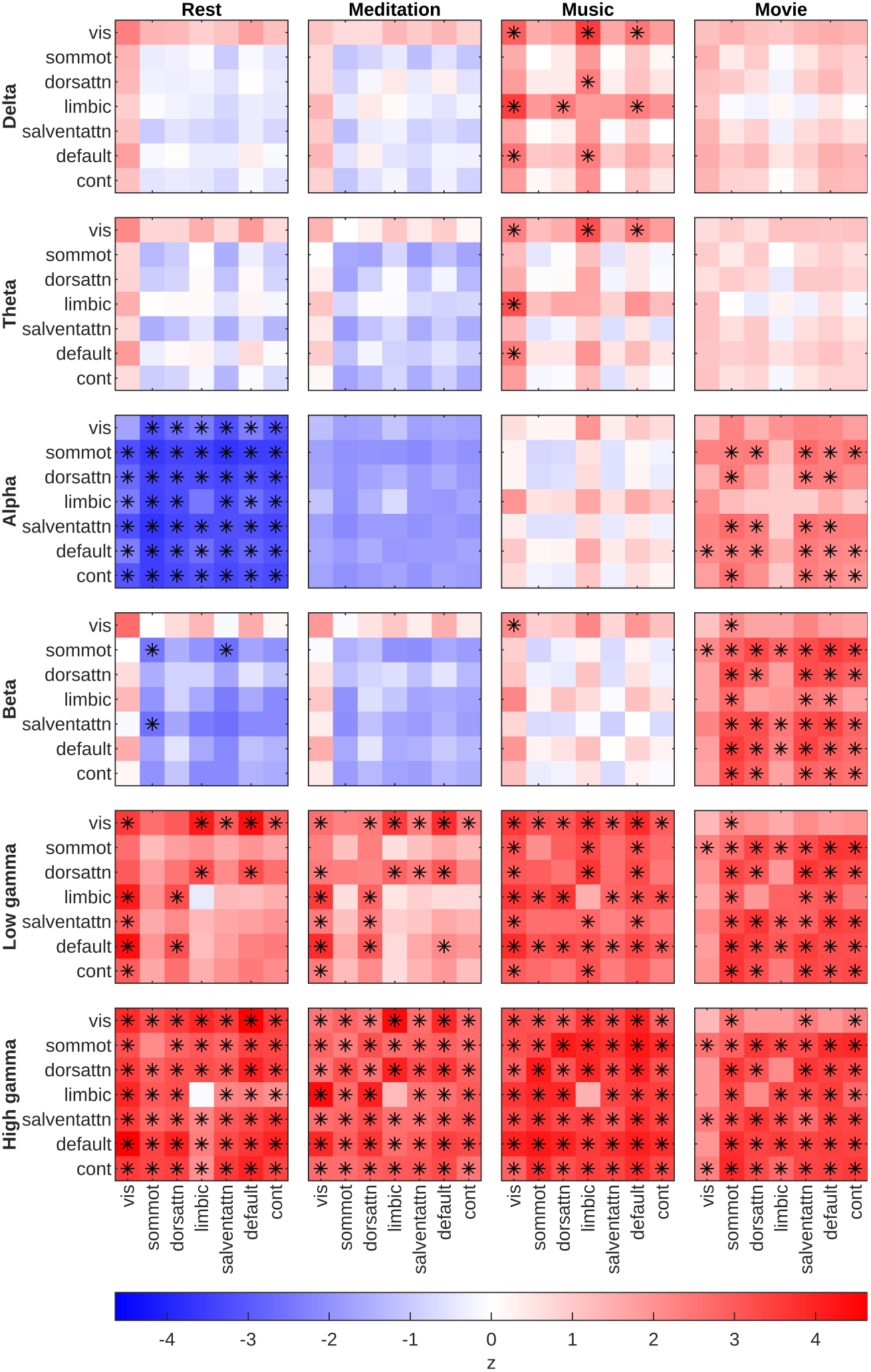
Difference in functional connectivity (FC) between psilocybin and baseline states for all conditions. Shades of red and blue indicate psilocybin-related increases and decreases in FC, respectively. (*p*<*0.05, non-parametric paired test, FDR-corrected).

**Fig. 4:**
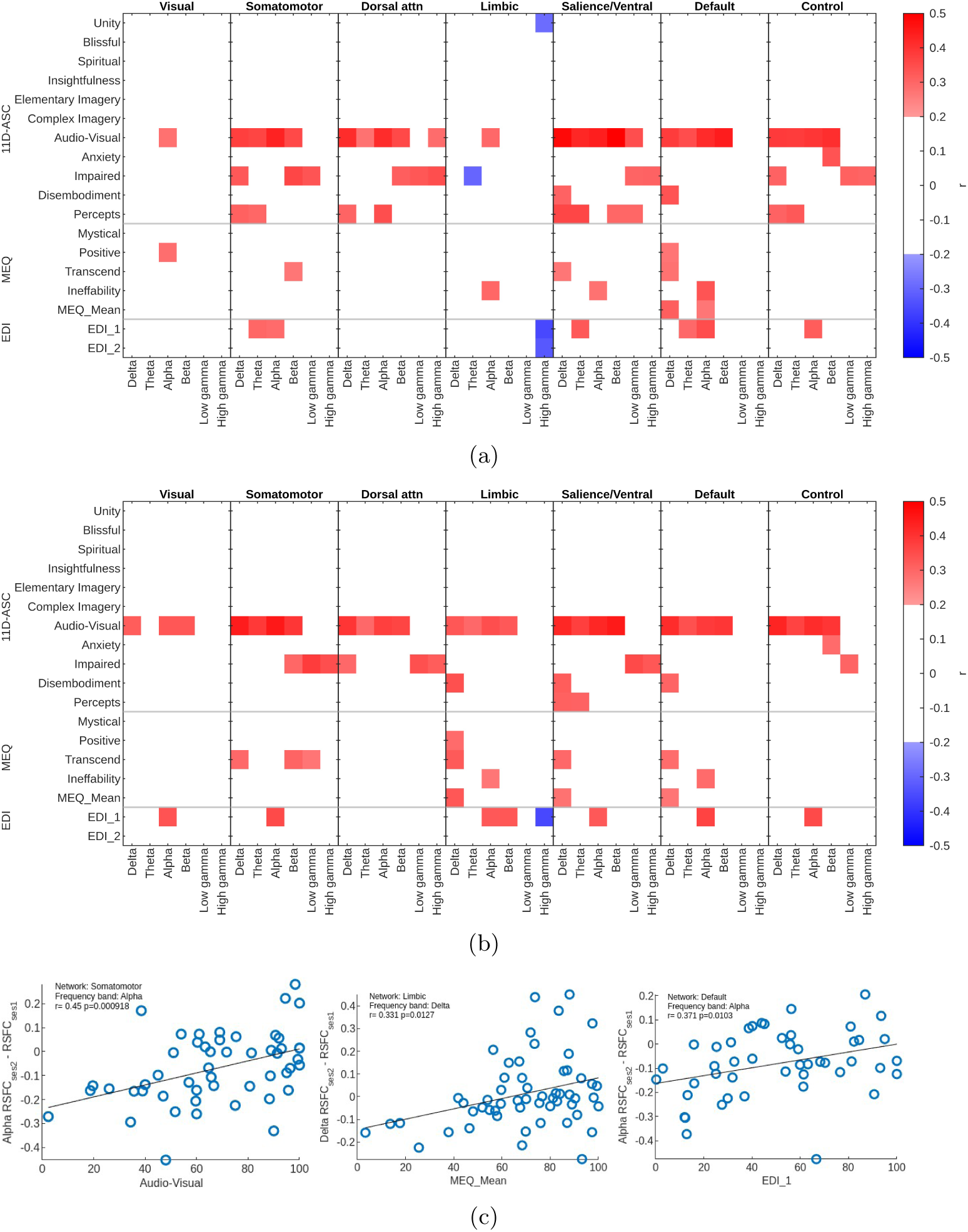
Spearman correlations between psilocybin-related changes in resting state functional connectivity (RSFC) and psychometric scales. (a) correlations within networks, and (b) correlations between each network and the rest of the brain. Only moderate and strong correlations (r*>*0.32, p*<*0.05, not FDR-corrected) are shown. Chosen psychometric scales include the 11-Dimensional Altered States of Consciousness (11D-ASC), Mystical Experience Questionnaire (MEQ), and Ego Dissolution Inventory (EDI), with EDI 1 and EDI 2 measured immediately before and after the EEG recording, respectively. (c) Scatter plots for the strongest correlations for each psychometric scale group, shown for clarity. The y-axis represents the change in RSFC between the psilocybin and baseline sessions for the corresponding network and frequency band: Somatomotor–alpha, limbic–delta, and default mode–alpha, from left to right.

#### 3.2.2 Context-dependent changes in alpha band functional connectivity

Across contexts, baseline-to-psilocybin alpha-band FC differences varied. Unlike rest, no within- or between-network alpha-band FC differences survived correction during meditation or music. (Figure 3, Figure S3, Figure S4). However under music, the psilocybin-related FC increase within the visual network was significant in all bands except alpha. Additionally, the limbic network showed notable increases in FC with the visual, DAN and DMN in the delta band; a trend not seen with any other contexts. Under meditation, none of the network pairs resulted in significant psilocybin-related FC changes in the low to mid frequency bands. In contrast to all eyes-closed conditions, movie-watching resulted in a broadband increase in FC across networks (Figure S5). Key networks associated with top-down control such as the DAN, S/VAN, DMN and control network (CN) showed significant increases in FC above 8 Hz. The SMN displayed a similar trend.

Comparing within context effects on FC while under baseline, the LME showed that while meditation did not alter FC from the resting state, music and movie-watching did (Figure 5). The condition of movie-watching produced the strongest decrease in FC across network pairs. Notably, although psilocybin alone reduced FC, the psilocybin × context interaction shifted FC in the opposite direction. Interestingly, no significant psilocybin × context interaction was detected for within-visual-network FC.

**Fig. 5:**
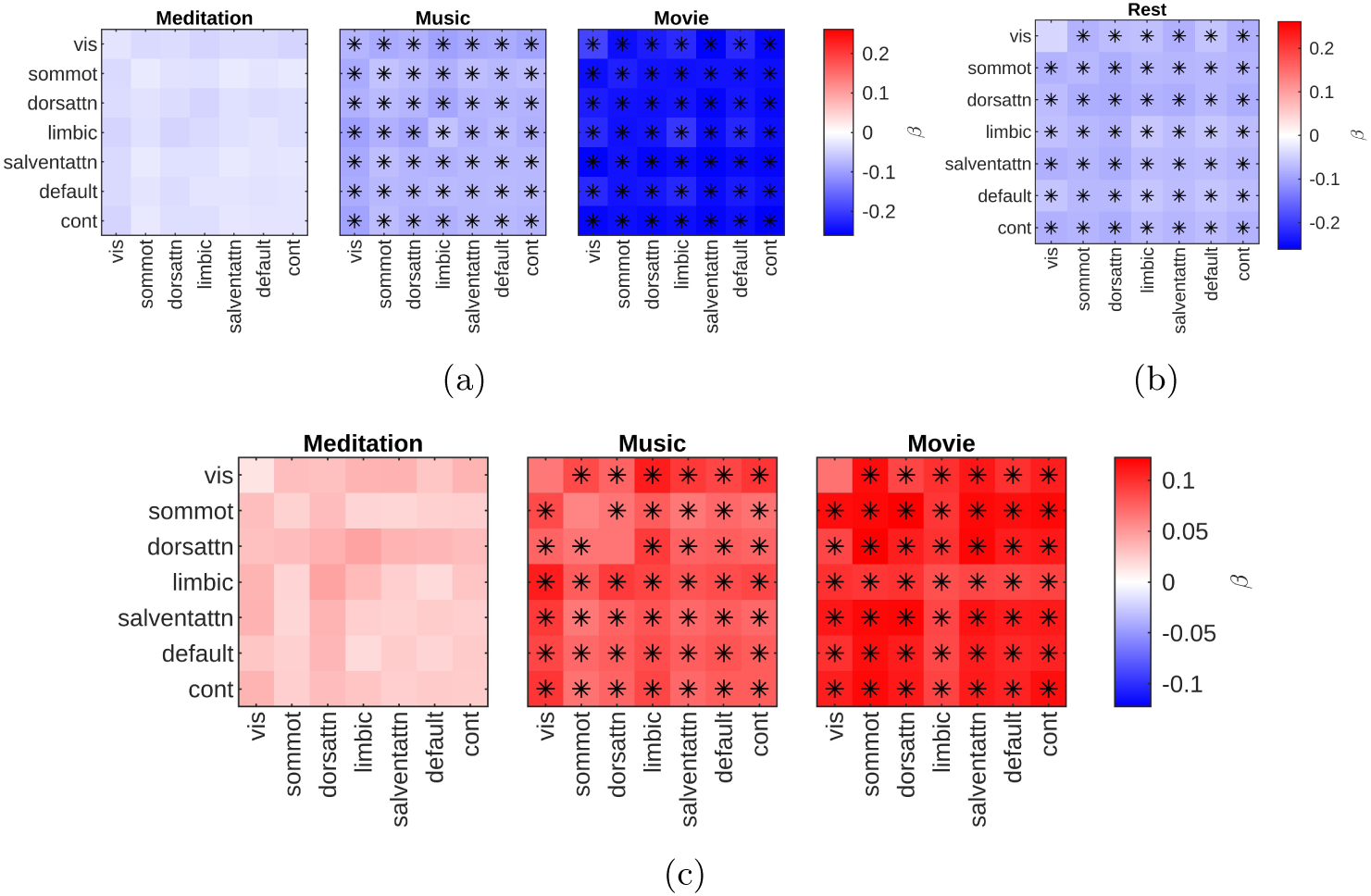
Drug, context and interaction effects (*β*) on alpha band functional connectivity (FC) within and between networks (*p*<*0.05, FDR-corrected). (a) Effect of naturalistic contexts on FC compared to rest during baseline session. (b) The drug effect on FC under rest. (c) The added interaction effect of drug and naturalistic contexts on FC (beyond the main effects of drug and condition). The color bar represents the *β* values, with blue and red indicating negative and positive effects, respectively.

#### 3.2.3 Network specific changes in the alpha band

To compare effects across networks, we summarised alpha-band FC as within-network connectivity and mean connectivity between each network and all other cortical networks. ( Figure S6). The psilocybin effect on mean RSFC was most pronounced for connections involving the the SMN (*β* = -0.0759, SE = 0.0207, p = 2.69 *×* 10^-4^), followed by the DAN, and S/VAN. Although significant changes were also observed in DMN, the magnitude of psilocybin-related effects was comparatively smaller. The psilocybin × music interaction was strongest for connections between the limbic network and other networks, followed by visual-to-other network interactions, with the DMN becoming more prominently involved. Under the movie context, the strongest effects were observed in the SMN and S/VAN.

Within individual networks, we observed that in the DMN: the FC decrease associated with the movie stimulus only (*β* = -0.2333, SE = 0.0218, p = 5.93 *×* 10^-24^) was stronger than the context-independent psilocybin effect (*β* = -0.0610, SE = 0.0215, p = 4.85 *×* 10^-3^). The interaction between psilocybin and music (*β* = 0.0818, SE = 0.0305, p = 7.55 *×* 10^-3^) as well as psilocybin and movie (*β* = 0.1033, SE = 0.0307, p = 8.48 *×* 10^-4^) opposed the negative psilocybin effect estimated during rest (Table 2). These trends were consistent in other key networks including the DAN (Table S4).

**Table 2:** Linear mixed-effects (LME) results for alpha band functional connectivity within the default mode network (*p*<*0.05, FDR-corrected).

| Within Default Mode network | | $\beta$ | SE | p |
| --- | --- | --- | --- | --- |
| Context independent | Drug | -0.0610 | 0.0215 | 4.85E-03* |
| Meditation | Context | -0.0297 | 0.0215 | 1.68E-01 |
|  | Drug x Context | 0.0207 | 0.0305 | 4.97E-01 |
| Music | Context | -0.0709 | 0.0215 | 1.08E-03* |
|  | Drug x Context | 0.0818 | 0.0305 | 7.55E-03* |
| Movie | Context | -0.2333 | 0.0218 | 5.93E-24* |
|  | Drug x Context | 0.1033 | 0.0307 | 8.48E-04* |

## 4 Discussion

The principal finding of this study is that key electrophysiological effects of psilocybin differed across contexts. At rest, psilocybin was associated with reduced low-frequency power and reductions in alpha-band connectivity across large-scale cortical networks implicated in top-down control and attentional regulation. Several of these effects were smaller or opposite in direction during music and movie viewing. These results suggest that the resting-state electrophysiological profile of psilocybin does not generalise uniformly across naturalistic conditions and therefore identifies context as a key moderator shaping the neural expression of the psychedelic state.

At rest, psilocybin was linked to an acute reduction in alpha-band functional connectivity across nearly all network pairs, except within visual and limbic networks. Although connectivity reductions were also observed in delta, theta, and beta bands, these effects were weaker and did not survive correction for multiple comparisons. Our findings extend previous MEG studies reporting reduced resting-state connectivity under psilocybin [4, 49] by systematically characterising frequency-specific FC within and between canonical functional networks (eg. DMN, S/VAN), showing that psilocybin-related connectivity changes are organised by both oscillatory frequency and functional network architecture.

The prominence of alpha-band effects is theoretically informative, as alpha activity has been associated with long-range cortical coordination, selective attention, and the regulation of cortical excitability [36]. Thus, reduced alpha-band connectivity under psilocybin may reflect a weakening of structured long-range coordination between networks involved in cognitive control, attention, and self-related processing. This finding may also have relevance for mechanistic models of antidepressant action, as increased delta-beta coherence has been associated with impaired top-down control in major depressive disorder (MDD) [40]. psilocybin-related reductions in alpha-band connectivity may reflect a loosening of rigid cortical coordination, providing a plausible electrophysiological pathway through which psilocybin could alter network dynamics implicated in depression. However, because the present study was conducted in healthy participants, and did not measure antidepressant response, this potential antidepressant mechanism remains speculative and must be further validated in clinical populations.

The networks most affected by psilocybin included the DMN, S/VAN, DAN, and frontoparietal control network, systems commonly associated with internally oriented cognition, salience processing and attentional control. These higher-order networks overlap with regions of high 5-HT_2*A*_ receptor density, positioning them as key targets of serotonergic psychedelic action [8, 21]. Consistent with prior fMRI studies [27, 57], psilocybin reduced DMN connectivity; however, DMN modulation was not the predominant effect. Rather, broader alpha-band connectivity reductions across higher-order networks suggest widespread reorganisation of oscillatory communication. This pattern is consistent with accounts of psychedelic states as desynchronised or less constrained [43, 57], including REBUS (Relaxed Beliefs Under Psychedelics) accounts proposing relaxed high-level priors and weakened top-down control over perception and cognition [13, 61, 62]. Such relaxation may permit novel interactions between sensory and associative systems to emerge [28, 66], potentially consistent with the relative preservation of visual and limbic connectivity observed here. These findings extend fMRI-based accounts by showing that large-scale psychedelic network reorganisation is expressed through specific oscillatory channels, with DMN disruption forming part of a broader alpha-band alteration across higher-order cortical systems.

Context-dependent connectivity changes indicated that psilocybin-related alpha-band effects varied with sensory and attentional context. Linear mixed-effects modelling showed that psilocybin alone reduced alpha-band connectivity, whereas the psilocybin *×* context interaction was associated with relative increases in connectivity. Meditation and music attenuated psilocybin-related alpha connectivity decreases, while movie watching reversed them and produced broader connectivity increases.

This pattern converged with the source spectral findings: psilocybin-related robust reductions at rest in theta, alpha, and beta bands, were progressively attenuated under naturalistic stimulation, with movie watching largely abolishing lower- to mid-frequency effects. Together, these results suggest that externally structured contexts constrain the neural expression of psilocybin, reducing the dominance of internally generated psychedelic dynamics. This interpretation aligns with recent MEG findings under LSD showing reduced psychedelic-related increases in neural complexity during external stimulation [41], and with fMRI evidence from the same cohort, in which psilocybin aligned brain activity more closely with the sensory context, making eyes-closed activity more similar to eyes-open activity [63]. Collectively, these findings support a competitive interaction between endogenous psychedelic dynamics and exogenous sensory input: under low-stimulation conditions, psilocybin-related dynamics may dominate cortical activity, whereas structured external input may organise neural activity around stimulus-driven processing.

Context effects varied across stimulus types. Music strongly modulated limbic connectivity, consistent with the role of limbic systems in emotion, affective auditory processing, and autobiographical experience [1, 37]. Given that music is central to psychedelic-assisted therapy and has been linked to mystical or insightful experiences and clinical improvement [5, 34], these findings support music as an active contextual modulator of the psychedelic state. In contrast, meditation produced weaker connectivity modulation than music or movie watching. Although meditation under psychedelics may alter neural dynamics and promote positive experiences with sustained or repeated practice [12, 50, 58], the brief meditation condition here may have been insufficient to produce a distinct neural state beyond rest.

Exploratory analyses suggested that AVS was related to lower-to-mid-frequency connectivity within and between several networks, indicating that the interplay between auditory and visual experience may depend on coordinated activity across distributed cortical systems under psilocybin. This association appeared more prominent to AVS, despite moderate correlations between AVS and related subjective dimensions, including elementary imagery, complex imagery, and changed meaning of percepts (r*>*0.5; see Figure S7b). This specificity may reflect a distinctive electrophysiological signature of audiovisual perceptual integration, although these results should be interpreted cautiously and require replication in independent samples.

The source spectral power findings provide a local oscillatory basis for the observed network-level reorganisation. Psilocybin induced widespread reductions in low-frequency source power, particularly in theta, alpha, and beta bands, alongside increases in gamma activity. These findings broadly align with previous M/EEG studies reporting reductions in oscillatory power and current source density under psilocybin and other serotonergic psychedelics highly comparable to psilocybin such as LSD, and DMT [4, 38, 43, 53, 67]. This convergence supports reduced low-mid frequency oscillatory activity, particularly in posterior and higher-order cortical regions, as a robust electrophysiological feature of serotonergic psychedelic action. Reduced alpha power may reflect weakened inhibitory constraints over cortical processing [33], while gamma increases may indicate heightened local cortical excitation and active information processing, although neither provides a direct measure of synaptic excitation or inhibition [38]. Together, these spectral changes are compatible with altered cortical excitability.

Reductions in theta and alpha activity are also compatible with accounts of relaxed top-down control. Theta activity, has been linked to higher attentional demands and cognitive load during processes such as memory encoding [17, 35], indicating that their reduction under psilocybin may therefore reflect weaker constraints on sensory and associative processing, consistent with predictive-processing accounts proposing relaxed high-level priors and increased bottom-up processing [13].

Despite broad convergence with prior work, some spectral findings differed from reports of broadband power reductions under psilocybin [43]. Frontal delta and gamma power increased, although gamma increases have also been reported under psilocybin [69]. The strongest gamma effects localised to medial cingulate and orbitofrontal regions (Table S1), rather than the lateral sites where muscle contamination dominates. However, source localisation cannot exclude non-neural contamination and given the susceptibility of gamma-band EEG to myogenic, ocular, movement, and line-noise artefacts, these findings should be interpreted cautiously. Differences from previous spectral findings may also reflect dose and recording timing. Psychedelic spectral effects appear to evolve dynamically, with early alpha decreases and delta/gamma increases preceding beta reductions [45], while microdosing studies have identified early theta reductions [18]. Thus, dose and temporal dynamics may contribute to discrepancies between the present study (19 mg oral) and earlier work using 2 mg intravenous psilocybin [43].

Beyond spectral power, psilocybin increased alpha peak frequency (APF) across most cortical regions and conditions. Higher APF has been linked to cognitive engagement, sensorimotor demands, and the timing of thalamo-cortical feedback [7, 42], and may reflect altered timing of alpha-mediated inhibitory cycles [36]. APF increases were particularly evident in parietal regions overlapping default mode and visual networks during rest, music, and movie conditions, suggesting that psilocybin alters alpha timing within the cortical systems most engaged by the ongoing state or task. APF was not significantly altered during meditation, possibly reflecting the brief duration of this condition, although this remains uncertain. APF therefore provides a complementary marker of psilocybin-related changes in alpha dynamics beyond power alone.

The visual network showed a distinct connectivity profile: psilocybin did not significantly reduce within-network alpha connectivity, but increased connectivity at other frequencies. This shift away from alpha dominance is consistent with reduced inhibitory gating within visual systems and aligns with prior serotonergic psychedelic studies reporting decreased alpha power and enhanced visual cortical connectivity [14, 39]. Together with alpha suppression and exploratory AVS-connectivity associations, this suggests frequency-specific reorganisation of communication between visual and higher-order systems, consistent with prior work linking psychedelic visual imagery to altered connectivity of visual cortex with higher-order systems [14, 39, 60]. Clinically, the context dependence of these effects reinforces the importance of setting in psychedelic-assisted interventions. ‘Grounding’, where directing attention externally reduces psilocybin-related network alteration, has been proposed as a means of reconnecting with external reality during therapy [57]. Contexts such as music listening and movie watching may act as active components of the psychedelic state, shaping how pharmacologically induced neural dynamics are expressed. Given the strong modulation of limbic connectivity by music, which is already central to psychedelic therapy protocols, these findings suggest that therapeutic environments may influence acute neural dynamics.

Certain limitations should be considered. Although connectivity was estimated at the source level to reduce the influence of field spread, which is particularly problematic for sensor-level connectivity analysis [54], source reconstruction does not fully eliminate signal leakage, and MSC remains sensitive to zero-lag correlations. While this does not preclude MSC as a biomarker for clinical studies, future work could validate the alpha-band findings using leakage-resistant measures such as imaginary coherence or phase-lag index. Template rather than individually digitised electrode positions may also have reduced spatial precision during source reconstruction. Although prior analyses found no monotonic effects of condition order [63], subtle temporal, expectancy, or carry-over effects cannot be excluded, particularly given the fixed-order design and absence of effective blinding. Finally, these findings reflect acute psilocybin effects in healthy participants and require validation in clinical populations and in relation to longer-term therapeutic outcomes.

## 5 Conclusion

Using the largest psilocybin-EEG cohort to date, this study presents the alpha band as a key frequency through which psilocybin reorganises cortical function. Critically, these effects were not fixed, but were shaped by setting. Externally structured contexts such as music listening and movie watching attenuated or reversed the psilocybin-related spectral and connectivity changes most pronounced at rest. Together, these findings suggest that psilocybin-related cortical dynamics reflect a balance between internally driven psychedelic processes and externally driven sensory input, with implications for how therapeutic settings are designed.

## Acknowledgements

We acknowledge all contributors involved in the design and acquisition of the original dataset and gratefully thank the volunteer participants whose involvement made this research possible. This research was supported by Monash eResearch capabilities, including high-performance computing resources (M3/MASSIVE).

VD was supported by a scholarship from the Monash AI Institute. AR received funding from the Australian Research Council (Grant Nos. DE170100128, DP200100757, FT250100563, and DP260104251), the Australian National Health and Medical Research Council (Investigator Grant No. 1194910), and a Turner Institute Strategic Grant, an internal Monash University initiative. AR is a CIFAR Global Scholar in the Brain, Mind & Consciousness Program and acknowledges support through the CIFAR Next-Generation Initiatives (Grant No. GS22-009) and Catalyst Grants (Grant Nos. CF-0224 and CF-0491).

VD was responsible for study design with input from LK, DS and AR. VD conducted all analyses; MB conducted data preprocessing. VD wrote the original draft of the manuscript; DS, LN, MB, AR, and LK provided critical input to data interpretation and reviewed and edited the manuscript.

All data reported are open access and available through OpenNeuro https://openneuro.org/datasets/ds006110; v1.2.0. Sensitive demographic information will be available upon signing a data access and confidentiality agreement (through the link available in the OpenNeuro data repository).

