## Supplementary material for "Context Shapes Alpha-Band Electrophysiological Signatures under Psilocybin"

### Context Shapes Alpha-Band Electrophysiological Signatures of Psilocybin-Induced Altered Consciousness

#### Supplementary material

##### Linear mixed effects model

The model is detailed as follows:

$$\text{MSC} = \beta_0 + \beta_{\text{Drug}} + \beta_{\text{Context}} + \beta_{\text{Drug}\times\text{Context}} + u_{\text{Subject}} + \epsilon \tag{1}$$

The main effect of drug reflects the overall change in FC between drug and baseline sessions, independent of context. The main effect of context represents FC differences across the contexts of meditation, music and movie, independent of drug. The drug × context interaction captures whether the effect of drug varies across context (for example, whether connectivity under the drug is more strongly reduced in meditation compared to music). A random gradient was included to account for the subject specific connectivity.

##### Reported Measures

Self-report measures were part of a broader assessment conducted before and longitudinally after psilocybin administration, with a subset included in the present analysis. The 11-Dimensional Altered States of Consciousness (11D-ASC) scale (42 items) [2, 4] was administered after the EEG recording to assess subjective alterations in consciousness. The Mystical Experience Questionnaire (MEQ30) [1] was collected at the same time to quantify mystical-type experiences. In addition, the Ego Dissolution Inventory (EDI) [3] was administered immediately before and after the EEG recording to assess transient experiences of ego dissolution. All three scales were collected during both the baseline and psilocybin sessions.

**Table S1**: Local maxima of psilocybin-related resting-state source power changes in MNI coordinates corresponding to Figure 2. For each frequency band, regions are ranked by absolute $t$-value, and the ten strongest effects are reported.

| Band | Region | t | MNI coordinates | | |
|---|---|---|---|---|---|
| **Delta** | | | | | |
| | superiorparietal L | -7.13 | -22.96 | -77.06 | 49.67 |
| | supramarginal R | -7.11 | 60.94 | -45.23 | 26.67 |
| | cuneus R | -7.07 | 3.85 | -61.70 | 14.42 |
| | cuneus L | -6.32 | -13.81 | -74.47 | 32.27 |
| | paracentral R | -6.22 | 2.60 | -35.67 | 54.08 |
| | pericalcarine R | -5.78 | 7.70 | -68.70 | 11.40 |
| | parstriangularis L | 5.58 | -44.92 | 38.74 | -9.19 |
| | pericalcarine L | -5.53 | -25.11 | -68.47 | 5.86 |
| | parsorbitalis L | 5.34 | -48.62 | 41.26 | -12.24 |
| | precuneus R | -5.19 | 16.03 | -73.09 | 36.86 |
| **Theta** | | | | | |
| | superiortemporal R | -9.54 | 63.80 | -47.56 | 14.14 |
| | inferiortemporal R | -8.18 | 63.58 | -52.21 | -13.96 |
| | middletemporal R | -8.12 | 46.76 | -40.80 | 6.85 |
| | cuneus R | -7.93 | 2.72 | -73.18 | 19.55 |
| | caudalanteriorcingulate L | -7.81 | -3.79 | 16.57 | 35.48 |
| | superiortemporal L | -7.56 | -54.70 | -24.21 | 11.54 |
| | lateraloccipital L | -7.20 | -39.13 | -69.54 | 4.66 |
| | lateraloccipital R | -6.89 | 38.70 | -74.66 | 7.61 |
| | parahippocampal L | -6.79 | -15.60 | -43.35 | -14.19 |
| | parahippocampal R | -6.74 | 29.69 | -42.08 | -7.84 |
| **Alpha** | | | | | |
| | precuneus L | -6.50 | -8.01 | -55.05 | 26.22 |
| | lingual R | -6.44 | 10.37 | -50.42 | -3.88 |
| | paracentral L | -6.35 | -2.94 | -35.47 | 50.78 |
| | pericalcarine L | -6.30 | -17.46 | -65.84 | 4.18 |
| | parahippocampal R | -5.74 | 16.53 | -29.75 | -10.71 |
| | cuneus L | -5.73 | -19.76 | -65.09 | 18.66 |
| | insula L | -5.67 | -31.30 | -26.15 | 13.52 |
| | lingual L | -5.64 | -21.95 | -61.45 | 3.10 |
| | precuneus R | -5.49 | 2.68 | -54.26 | 27.14 |
| | fusiform R | -5.40 | 43.28 | -54.99 | -23.63 |
| **Beta** | | | | | |
| | precuneus L | -7.39 | -11.74 | -47.90 | 33.23 |
| | inferiorparietal L | -7.13 | -25.16 | -64.94 | 29.77 |
| | isthmuscingulate R | -7.07 | 9.31 | -37.93 | 3.89 |
| | lingual R | -7.07 | 21.90 | -63.08 | 3.91 |
| | isthmuscingulate L | -6.78 | -10.64 | -48.48 | 31.23 |
| | pericalcarine R | -6.74 | 5.02 | -66.16 | 11.88 |
| | precuneus R | -6.17 | 14.51 | -64.68 | 23.69 |
| | lingual L | -6.14 | -9.78 | -67.96 | 4.68 |
| | caudalanteriorcingulate R | -6.11 | 1.42 | 15.87 | 27.14 |
| | cuneus L | -5.99 | -17.86 | -80.48 | 38.71 |
| **Low gamma** | | | | | |
| | medialorbitofrontal R | 6.01 | 8.91 | 24.90 | -24.74 |
| | medialorbitofrontal L | 5.76 | 0.32 | 20.48 | -26.42 |
| | lateralorbitofrontal R | 5.75 | 13.76 | 21.22 | -16.41 |
| | lateralorbitofrontal L | 5.64 | -9.41 | 13.01 | -24.60 |
| | superiortemporal R | 5.50 | 21.46 | 12.67 | -33.87 |
| | entorhinal R | 5.45 | 15.53 | 1.58 | -26.72 |
| | superiortemporal L | 5.43 | -32.64 | 23.45 | -36.55 |
| | parsorbitalis L | 5.31 | -30.65 | 26.51 | -5.34 |
| | rostralmiddlefrontal L | 4.92 | -35.26 | 37.48 | -13.86 |
| | caudalanteriorcingulate L | 4.66 | -2.70 | 30.22 | 12.59 |
| **High gamma** | | | | | |
| | rostralanteriorcingulate L | 5.85 | -1.75 | 18.49 | -6.47 |
| | parahippocampal L | 5.48 | -13.83 | -30.92 | -13.07 |
| | caudalanteriorcingulate L | 5.44 | -2.70 | 30.22 | 12.59 |
| | medialorbitofrontal R | 5.38 | 1.92 | 26.96 | -22.46 |
| | rostralanteriorcingulate R | 5.27 | 1.19 | 10.95 | -4.18 |
| | isthmuscingulate L | 5.23 | -13.99 | -53.17 | 3.56 |
| | lateralorbitofrontal R | 5.22 | 13.76 | 21.22 | -16.41 |
| | isthmuscingulate R | 5.18 | 13.13 | -40.88 | -2.81 |
| | medialorbitofrontal L | 5.17 | 0.15 | 27.21 | -27.89 |
| | superiorfrontal R | 5.01 | 0.56 | 41.92 | -14.89 |

L: Left, R: Right

**Table S2**: Local maxima of psilocybin-related source power changes in MNI coordinates corresponding to Figure 2 during meditation. For each frequency band, regions are ranked by absolute $t$-value, and the ten strongest effects are reported.

| **Band** | **Region** | **t** | MNI coordinates | | |
|---|---|---|---|---|---|
| **Delta** | | | | | |
| | cuneus R | -7.52 | 2.9 | -80.45 | 33.91 |
| | lingual L | -7.34 | -12.53 | -64.75 | 5.54 |
| | fusiform L | -7.17 | -43.86 | -57.72 | -9.55 |
| | pericalcarine L | -7.07 | -25.11 | -68.47 | 5.86 |
| | cuneus L | -7.03 | -13.89 | -71.20 | 25.53 |
| | superiorparietal L | -6.90 | -26.39 | -73.19 | 28.35 |
| | precuneus R | -6.87 | 16.22 | -72.10 | 31.83 |
| | posteriorcingulate L | -6.85 | -0.20 | -27.81 | 44.06 |
| | lingual R | -6.82 | 4.59 | -68.65 | 12.29 |
| | superiorparietal R | -5.96 | 32.01 | -76.95 | 22.81 |
| **Theta** | | | | | |
| | parahippocampal L | -9.57 | -24.77 | -27.64 | -28.87 |
| | inferiortemporal L | -8.95 | -41.89 | -33.93 | -28.60 |
| | lateralorbitofrontal R | -8.51 | 22.96 | 0.00 | -15.33 |
| | parahippocampal R | -8.50 | 21.61 | -38.37 | -14.72 |
| | lingual L | -8.36 | -19.62 | -67.09 | 4.00 |
| | fusiform L | -8.29 | -30.15 | -62.94 | -19.04 |
| | superiorparietal R | -8.24 | 29.89 | -62.83 | 58.39 |
| | inferiortemporal R | -7.90 | 62.74 | -58.28 | -6.00 |
| | lateraloccipital L | -7.81 | -37.81 | -66.16 | 14.61 |
| | precuneus R | -7.67 | 3.40 | -51.45 | 37.33 |
| **Alpha** | | | | | |
| | caudalanteriorcingulate L | -5.65 | -4.95 | 21.70 | 31.49 |
| | caudalanteriorcingulate R | -4.84 | 2.23 | 6.12 | 28.66 |
| | fusiform R | -4.78 | 34.75 | -67.08 | -19.47 |
| | middletemporal L | -4.63 | -63.95 | -40.77 | -16.79 |
| | cuneus L | -4.43 | -13.46 | -74.78 | 27.41 |
| | inferiortemporal R | -4.33 | 46.97 | -33.05 | -23.14 |
| | lingual L | -4.21 | -12.53 | -64.75 | 5.54 |
| | rostralmiddlefrontal L | 4.00 | -36.08 | 61.24 | -4.16 |
| | posteriorcingulate L | -3.97 | -3.81 | -29.48 | 26.71 |
| | parsopercularis L | -3.91 | -33.34 | 3.44 | 11.17 |
| **Beta** | | | | | |
| | isthmuscingulate R | -7.922 | 7.09 | -39.64 | 5.31 |
| | caudalanteriorcingulate L | -5.298 | -8.56 | 19.31 | 33.35 |
| | lingual R | -5.282 | 9.98 | -62.84 | 6.80 |
| | precuneus L | -5.169 | -13.58 | -65.64 | 28.95 |
| | isthmuscingulate L | -4.991 | -8.29 | -48.17 | 29.12 |
| | inferiorparietal R | -4.896 | 43.95 | -49.73 | 50.58 |
| | caudalanteriorcingulate R | -4.586 | 3.42 | 22.34 | 34.77 |
| | lingual L | -4.392 | -20.04 | -64.97 | 3.04 |
| | pericalcarine R | -4.076 | 5.02 | -66.16 | 11.88 |
| | inferiorparietal L | -3.924 | -23.41 | -64.29 | 38.49 |
| **Low gamma** | | | | | |
| | inferiortemporal R | 4.54 | 19.53 | -1.03 | -48.31 |
| | middletemporal L | 4.50 | -46.68 | 7.39 | -45.98 |
| | inferiortemporal L | 4.50 | -38.05 | 6.43 | -47.01 |
| | lateralorbitofrontal R | 4.45 | 10.10 | 34.77 | -24.82 |
| | parsopercularis L | 4.31 | -42.35 | 9.65 | -0.71 |
| | entorhinal R | 4.24 | 18.68 | 3.53 | -21.68 |
| | superiortemporal R | 4.21 | 26.81 | 16.03 | -30.83 |
| | isthmuscingulate L | 4.21 | -6.54 | -42.99 | 0.36 |
| | precuneus R | 4.11 | 11.64 | -56.17 | 7.79 |
| | isthmuscingulate R | 4.04 | 12.72 | -56.13 | 6.48 |
| **High gamma** | | | | | |
| | precuneus R | 5.31 | 11.35 | -55.29 | 13.24 |
| | isthmuscingulate R | 5.28 | 16.24 | -50.51 | 3.46 |
| | cuneus R | 4.91 | 18.72 | -71.70 | 22.09 |
| | pericalcarine R | 4.72 | 21.67 | -75.45 | 3.52 |
| | fusiform R | 4.56 | 25.60 | -76.75 | -7.16 |
| | parahippocampal R | 4.40 | 14.68 | -38.40 | -7.41 |
| | inferiortemporal L | 4.36 | -56.03 | -13.58 | -40.99 |
| | isthmuscingulate L | 4.34 | -8.28 | -51.83 | 1.80 |
| | precuneus L | 4.29 | -1.52 | -63.82 | 13.57 |
| | pericalcarine L | 4.28 | -4.17 | -87.64 | 0.93 |

**Table S3**: Local maxima of psilocybin-related source power changes in MNI coordinates corresponding to Figure 2 during music. For each frequency band, regions are ranked by absolute *t*-value, and the ten strongest effects are reported.

| Band | Region | t | MNI coordinates | | |
|---|---|---|---|---|---|
| **Delta** | | | | | |
| | cuneus R | -6.81 | 20.28 | -70.15 | 23.16 |
| | precuneus R | -6.77 | 20.44 | -67.66 | 22.98 |
| | superiorparietal R | -5.15 | 30.59 | -81.72 | 30.21 |
| | superiorparietal L | -4.44 | -26.39 | -73.19 | 28.35 |
| | cuneus L | -4.35 | -17.86 | -80.48 | 38.71 |
| | lateralorbitofrontal R | 3.93 | 30.07 | 61.20 | -14.88 |
| | parsorbitalis R | 3.50 | 47.05 | 45.47 | -17.16 |
| | parsorbitalis L | 3.31 | -50.46 | 37.90 | -14.76 |
| | rostralmiddlefrontal R | 3.30 | 49.82 | 45.23 | -8.52 |
| | superiorfrontal R | 3.25 | 2.33 | 47.27 | -0.96 |
| **Theta** | | | | | |
| | middletemporal R | -7.97 | 51.29 | -50.57 | -0.73 |
| | lateraloccipital R | -7.97 | 47.28 | -69.83 | -9.12 |
| | fusiform R | -7.73 | 21.79 | -66.62 | -11.53 |
| | inferiorparietal R | -7.44 | 30.07 | -80.72 | 18.32 |
| | superiorparietal L | -7.35 | -19.99 | -66.06 | 50.72 |
| | precuneus R | -7.15 | 11.19 | -49.84 | 41.26 |
| | isthmuscingulate R | -7.05 | 8.10 | -39.49 | 35.69 |
| | lingual L | -6.93 | -7.24 | -66.00 | 5.34 |
| | parahippocampal R | -6.87 | 34.40 | -32.06 | -17.05 |
| | supramarginal L | -6.72 | -32.68 | -29.88 | 19.50 |
| **Alpha** | | | | | |
| | caudalanteriorcingulate L | -5.46 | -1.86 | 20.03 | 36.26 |
| | precentral L | -4.51 | -36.41 | 4.82 | 30.74 |
| | rostralmiddlefrontal R | 4.47 | 38.73 | 51.31 | -4.86 |
| | rostralmiddlefrontal L | 4.46 | -37.71 | 50.80 | -4.49 |
| | parstriangularis R | 4.39 | 44.55 | 40.16 | -5.12 |
| | posteriorcingulate R | -4.37 | 2.89 | -5.96 | 30.43 |
| | superiorparietal R | -4.34 | 38.67 | -55.03 | 61.20 |
| | cuneus L | -4.14 | -17.86 | -80.48 | 38.71 |
| | postcentral R | -4.06 | 50.42 | -22.00 | 39.87 |
| | insula R | -3.81 | 41.72 | -14.88 | 13.52 |
| **Beta** | | | | | |
| | superiorfrontal R | -6.39 | 1.90 | 29.94 | 47.27 |
| | parsopercularis R | -6.34 | 35.31 | 8.70 | 26.88 |
| | caudalanteriorcingulate L | -5.91 | -1.52 | 20.50 | 37.74 |
| | caudalanteriorcingulate R | -5.68 | 2.68 | 7.67 | 28.22 |
| | superiorfrontal L | -5.21 | -7.57 | 22.89 | 43.23 |
| | isthmuscingulate L | -5.09 | -3.97 | -33.24 | 25.56 |
| | caudalmiddlefrontal R | -4.96 | 34.41 | 9.39 | 28.09 |
| | superiorparietal L | -4.91 | -22.14 | -62.25 | 35.30 |
| | isthmuscingulate R | -4.55 | 4.04 | -32.30 | 26.60 |
| | posteriorcingulate R | -4.51 | 3.22 | -2.18 | 30.21 |
| **Low gamma** | | | | | |
| | rostralanteriorcingulate L | 5.53 | -5.23 | 37.26 | -3.83 |
| | lingual R | 4.88 | 6.00 | -83.35 | -17.46 |
| | rostralanteriorcingulate R | 4.73 | 2.42 | 44.43 | 5.49 |
| | cuneus R | 4.69 | 4.48 | -92.27 | 0.63 |
| | medialorbitofrontal R | 4.56 | 1.92 | 26.96 | -22.46 |
| | superiorfrontal R | 4.55 | 2.22 | 45.33 | -14.09 |
| | medialorbitofrontal L | 4.49 | 0.15 | 27.21 | -27.89 |
| | pericalcarine R | 4.26 | 14.75 | -94.80 | -4.69 |
| | lateralorbitofrontal R | 4.22 | 28.72 | 16.46 | -27.54 |
| | caudalanteriorcingulate L | 4.16 | -2.70 | 30.22 | 12.59 |
| **High gamma** | | | | | |
| | cuneus R | 5.68 | 15.71 | -59.01 | 18.13 |
| | pericalcarine R | 5.36 | 21.67 | -75.45 | 3.52 |
| | precuneus R | 5.26 | 3.16 | -54.55 | 9.15 |
| | cuneus L | 5.07 | 0.49 | -68.10 | 14.88 |
| | isthmuscingulate R | 5.04 | 5.97 | -43.15 | 6.53 |
| | precuneus L | 4.96 | -9.19 | -58.54 | 13.04 |
| | lingual R | 4.78 | 19.05 | -73.61 | -17.44 |
| | caudalanteriorcingulate L | 4.71 | -2.70 | 30.22 | 12.59 |
| | caudalanteriorcingulate R | 4.66 | 5.62 | 31.10 | 11.35 |
| | isthmuscingulate L | 4.65 | -1.65 | -50.62 | 12.71 |

L: Left, R: Right

**Table S4**: Linear mixed-effects (LME) results for alpha band functional connectivity within the dorsal attention network (*p<0.05, FDR-corrected).

| Within Dorsal Attention network | | $\beta$ | SE | p |
|---|---|---|---|---|
| Context independent | Drug | -0.0847 | 0.0236 | 3.71E-04* |
| Meditation | Context | -0.0364 | 0.0236 | 1.24E-01 |
| | Drug x Context | 0.0379 | 0.0334 | 2.56E-01 |
| Music | Context | -0.0761 | 0.0236 | 1.37E-03* |
| | Drug x Context | 0.0668 | 0.0334 | 4.61E-02 |
| Movie | Context | -0.245 | 0.0239 | 2.66E-22* |
| | Drug x Context | 0.108 | 0.0337 | 1.44E-03* |

**Table S5**: Linear mixed-effects (LME) results for alpha-band functional connectivity within the visual network (*p<0.05, FDR-corrected).

| Within visual network | | $\beta$ | SE | p |
|---|---|---|---|---|
| Context independent | Drug | -0.0410 | 0.0244 | 0.0936 |
| Meditation | Context | -0.0282 | 0.0244 | 0.248 |
| | Drug × Context | 0.0133 | 0.0345 | 0.700 |
| Music | Context | -0.0760 | 0.0244 | 0.00193* |
| | Drug × Context | 0.0643 | 0.0345 | 0.0627 |
| Movie | Context | -0.191 | 0.0246 | 6.81E-14* |
| | Drug × Context | 0.0673 | 0.0348 | 0.0537 |

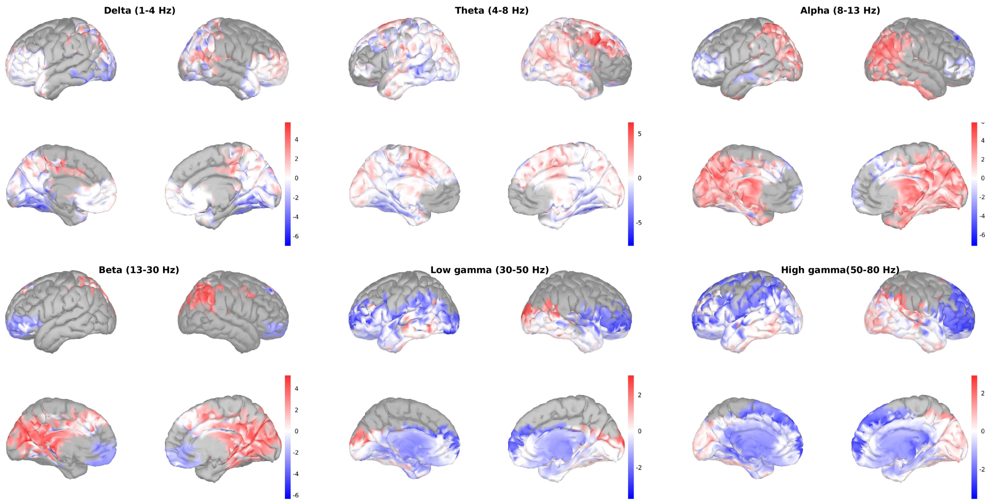


(a)

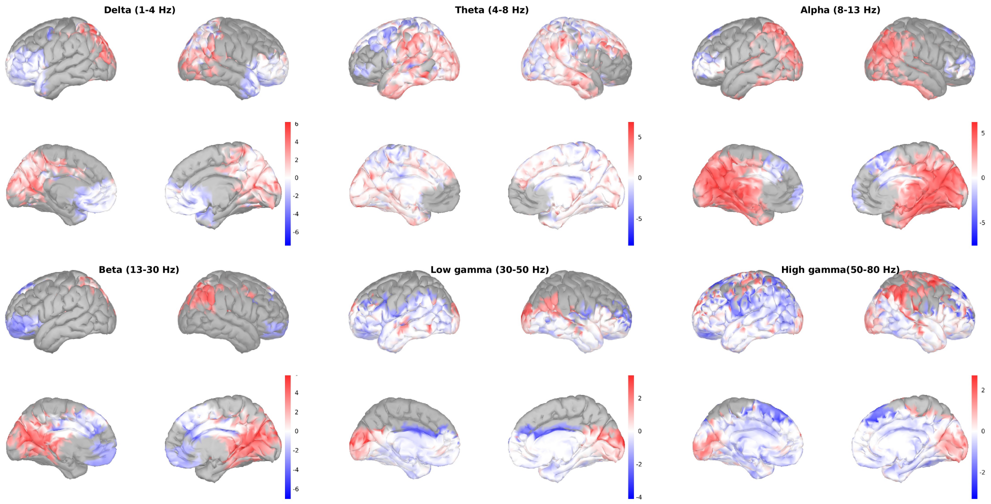


(b)

**Fig. S1**: Condition-dependent modulation of psilocybin-induced changes in source spectral power. Statistical parametric maps showing significant condition contrasts in psilocybin-induced source spectral power changes relative to rest. Contrasts are shown for (a) music versus rest and (b) meditation versus rest. Positive t-values indicate regions where the psilocybin-induced change in power was greater in the music or meditation condition than in the resting-state condition. Where the original psilocybin-induced effect reflected a reduction in power, positive t-values therefore indicate a weaker psilocybin-induced decrease under music or meditation compared with rest. Only significant effects are shown ($p<0.05$, corrected

).

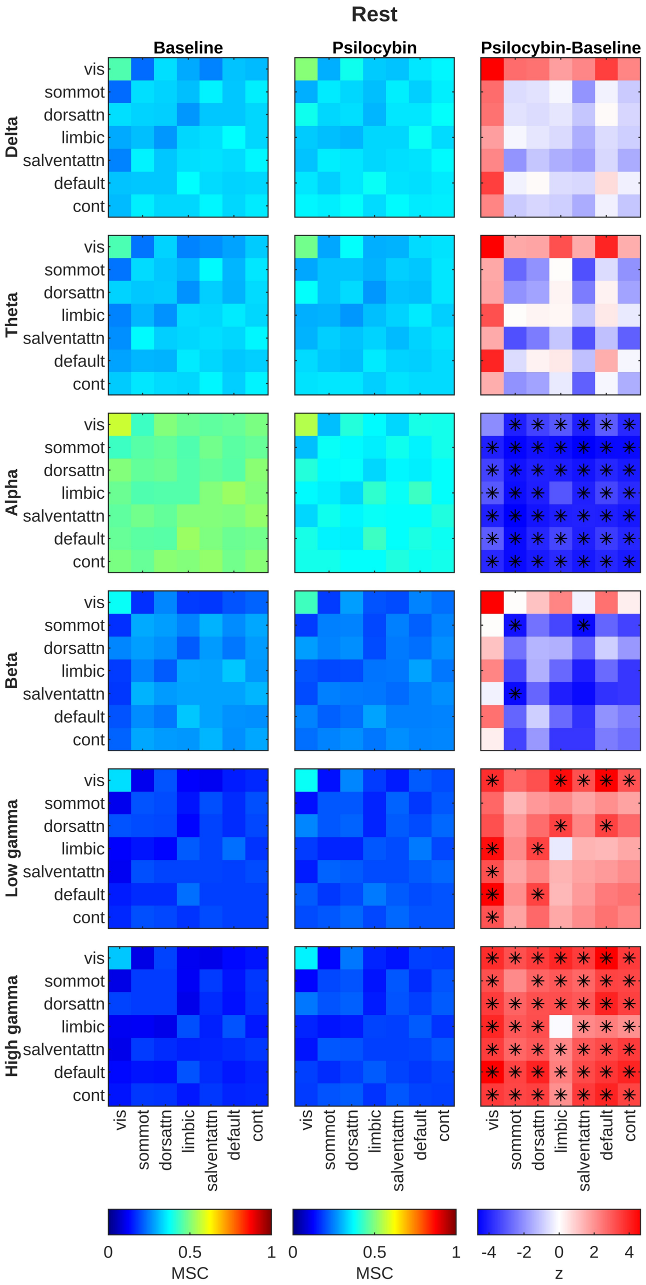


**Fig. S2**: Within and between network connectivity in the baseline and psilocybin states, followed by the difference between psilocybin and baseline states under rest (*p<0.05, non-parametric paired test, FDR-corrected).

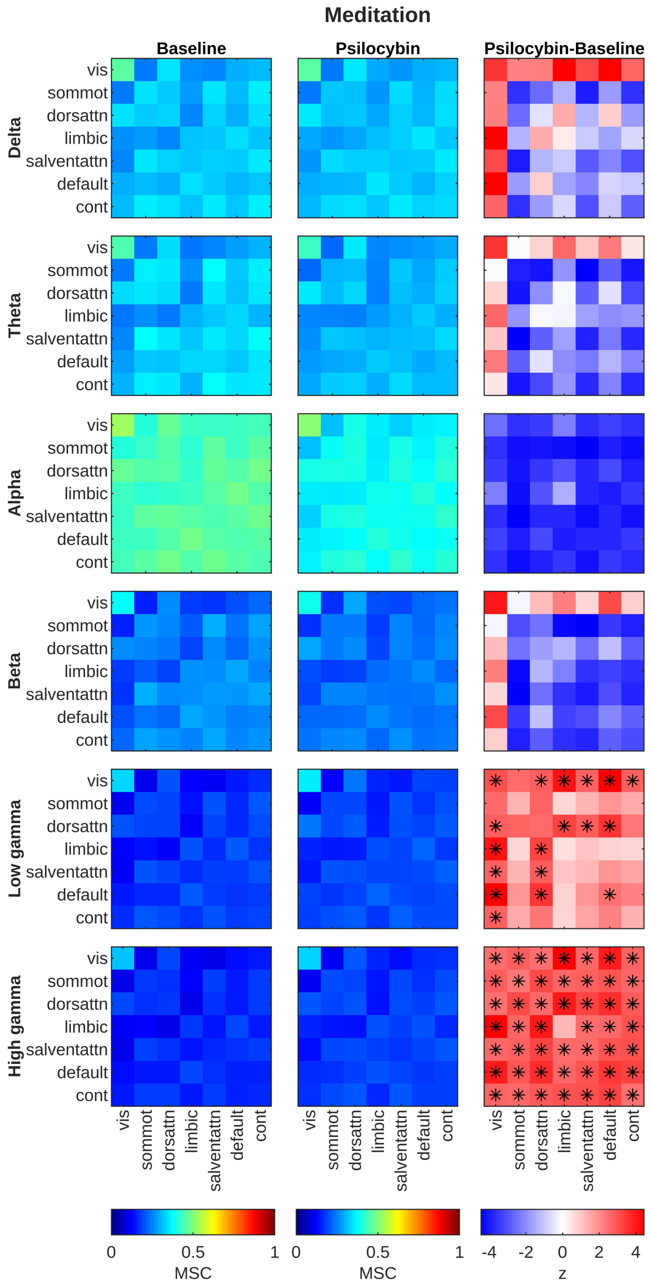


**Fig. S3**: Within and between network connectivities in the baseline and psilocybin states, followed by the difference between psilocybin and baseline states under meditation (*p<0.05, non-parametric paired test, FDR-corrected).

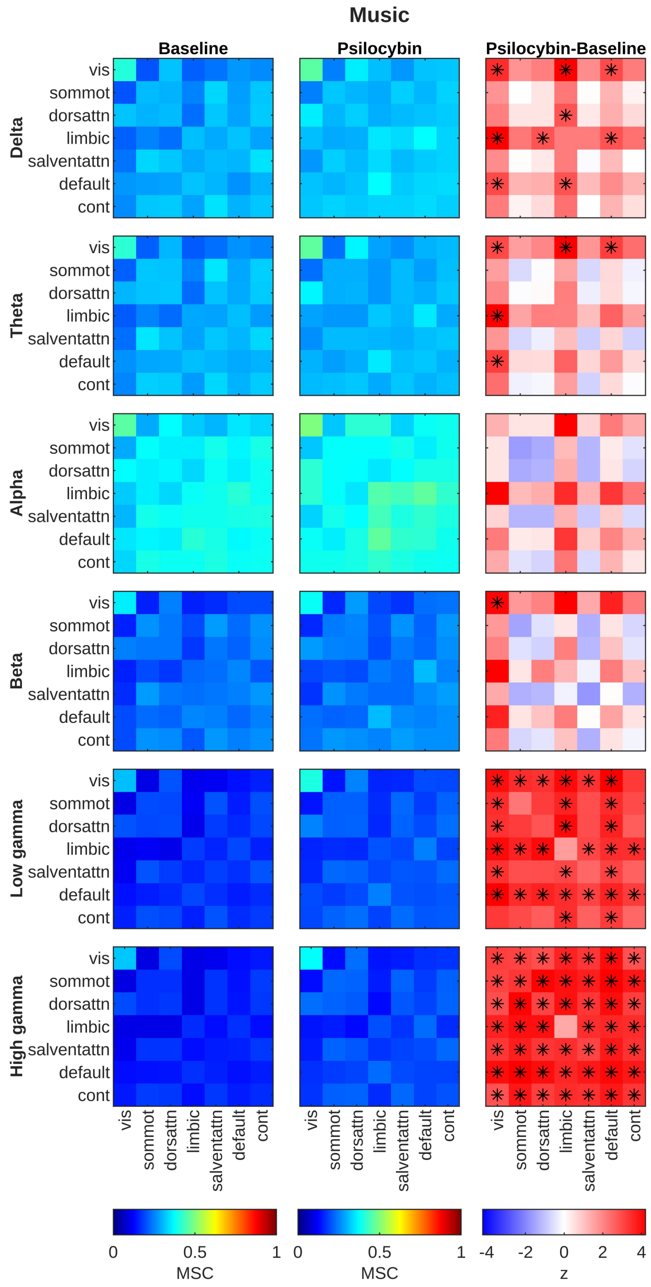


**Fig. S4**: Within and between network connectivities in the baseline and psilocybin states, followed by the difference between psilocybin and baseline states under music (*p<0.05, non-parametric paired test, FDR-corrected).

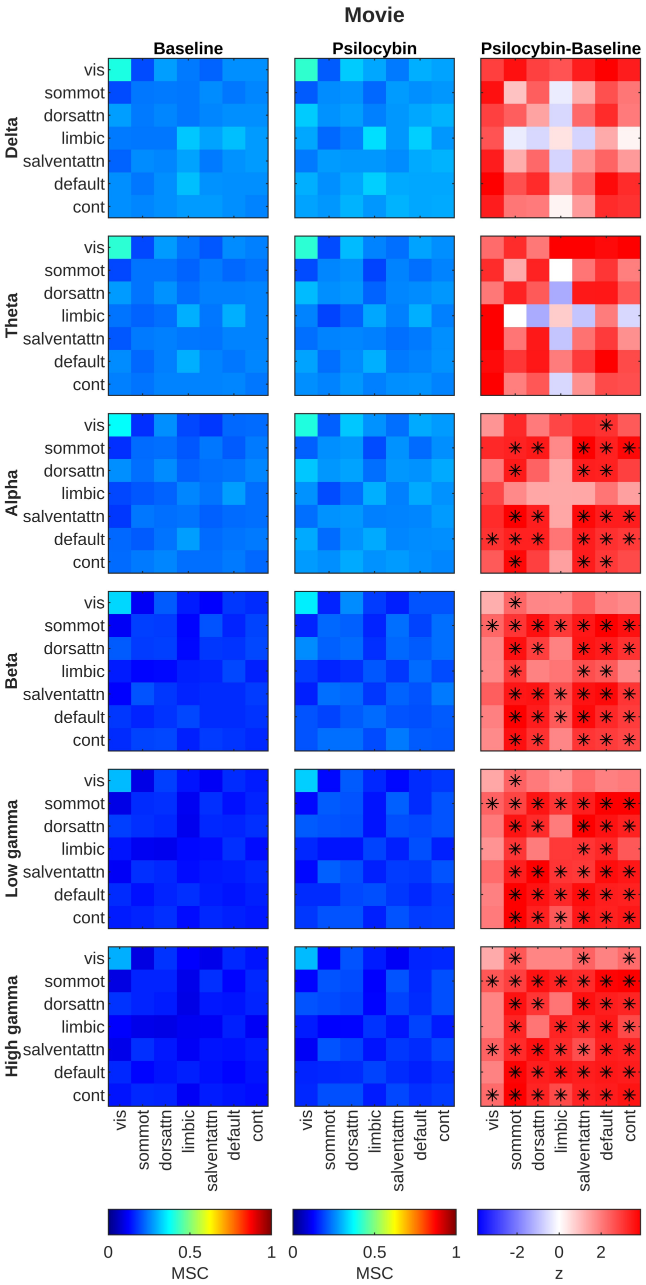


**Fig. S5**: Within and between network connectivities in the baseline and psilocybin states, followed by the difference between psilocybin and baseline states under movie-watching (*p<0.05, non-parametric paired test, FDR-corrected).

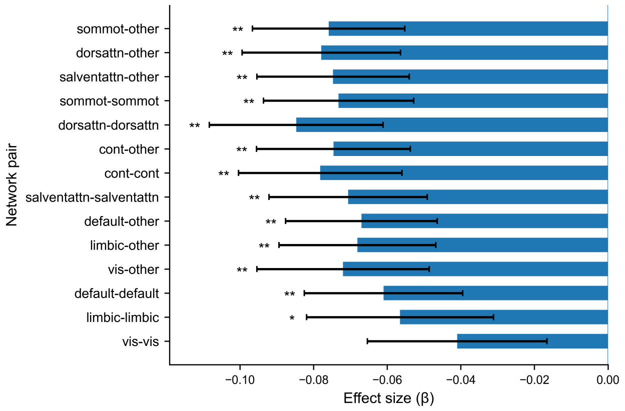


(a)

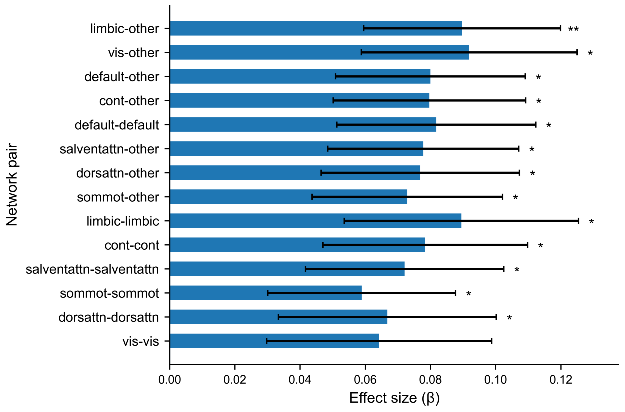


(b)

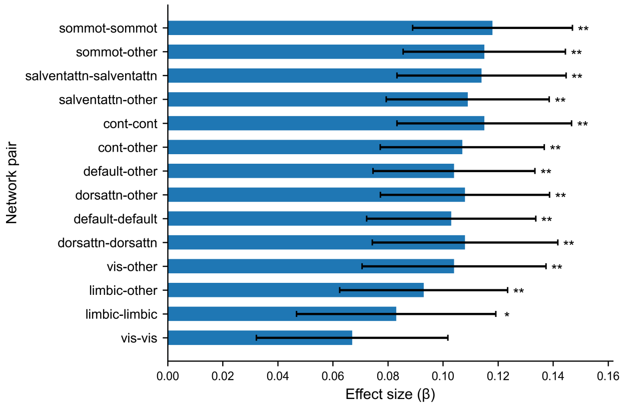


(c)

**Fig. S6**: Linear mixed model (LMM) effects on mean within- and between-network functional connectivity during resting state, ordered by probability (p) (*$p<0.05$, **$p<0.005$, FDR-corrected). (a) Drug effects, (b) added interaction effects of drug and music, and (c) added interaction effects of drug and movie.

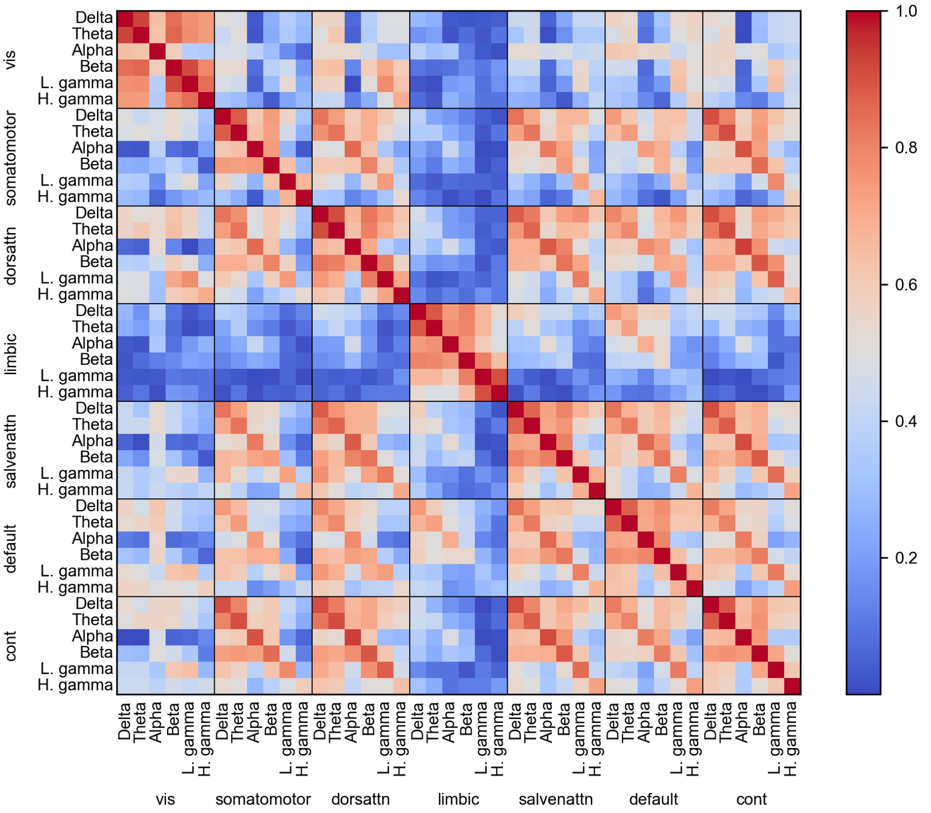


(a)

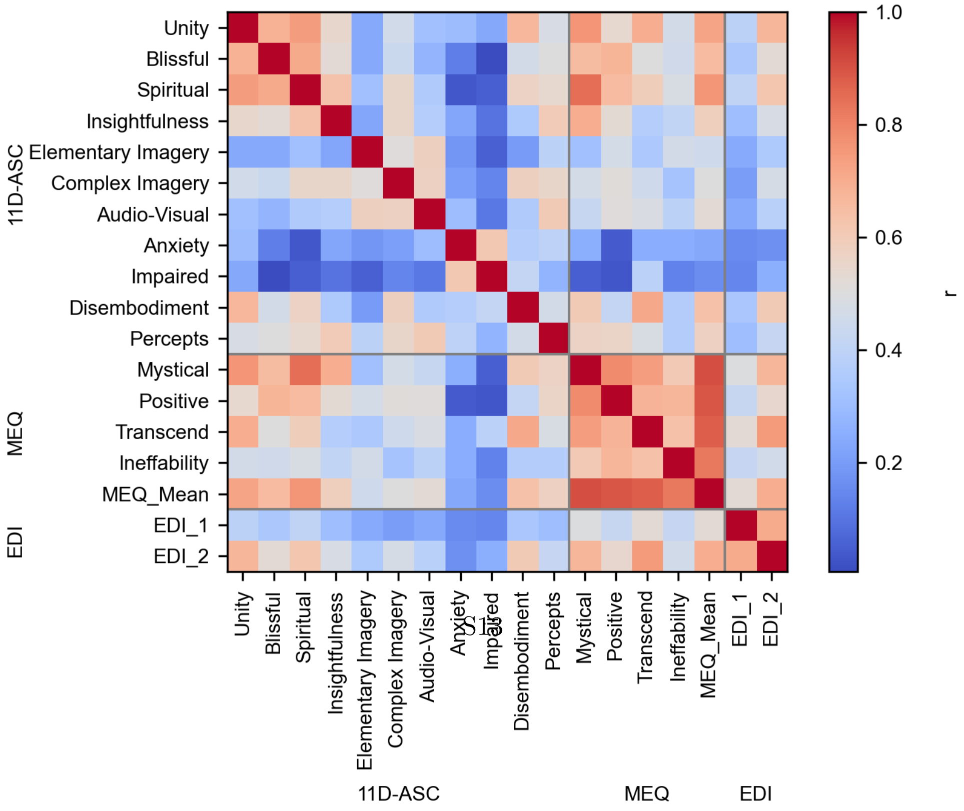


(b)

**Fig. S7**: Pearson correlations between (a) mean psilocybin-related changes in resting state functional connectivity (RSFC) within networks across frequency bands and (b) psychometric scales. Scales include 11-Dimensional Altered States of Consciousness (11D-ASC), Mystical Experience Questionnaire (MEQ), and Ego Dissolution Inventory (EDI), with EDI_1 and EDI_2 measured immediately before and after the EEG recording, respectively.
